# Cyclic transitions between higher order motifs underlie sustained activity in asynchronous sparse recurrent networks

**DOI:** 10.1101/777219

**Authors:** Kyle Bojanek, Yuqing Zhu, Jason MacLean

## Abstract

Many studies have demonstrated the prominence of higher-order patterns in excitatory synaptic connectivity as well as activity in neocortex. Surveyed as a whole, these results suggest that there may be an essential role for higher-order patterns in neocortical function. In order to stably propagate signal within and between regions of neocortex, the most basic - yet nontrivial - function which neocortical circuitry must satisfy is the ability to maintain stable spiking activity over time. Here we algorithmically construct spiking neural network models comprised of 5000 neurons using topological statistics from neocortex and a set of objective functions that identify networks which produce naturalistic low-rate, asynchronous, and critical activity. We find that the same network topology can exhibit either sustained activity under one set of initial membrane voltages or truncated activity under a different set. Yet these two outcomes are not readily differentiated by rate or criticality. By summarizing the statistical dependencies in the pairwise activity of neurons as directed weighted functional networks, we examined the transient manifestations of higher-order motifs in the functional networks across time. We find that stereotyped low variance cyclic transitions between three isomorphic triangle motifs, quantified as a Markov process, are required for sustained activity. If the network fails to engage the dynamical regime characterized by a recurring stable pattern of motif dominance, spiking activity ceased. Motif cycling generalized across manipulations of synaptic weights and across topologies, demonstrating the robustness of this dynamical regime for sustained spiking in critical asynchronous network activity. Our results point to the necessity of higher-order patterns amongst excitatory connections for sustaining activity in sparse recurrent networks. They also provide a possible explanation as to why such excitatory synaptic connectivity and activity patterns have been prominently reported in neocortex.

**Author summary:** Here we address two questions. First, it remains unclear how activity propagates stably through a network since neurons are leaky and connectivity is sparse and weak. Second, higher order patterns abound in neocortex, hinting at potential functional relevance for their presence. Several lines of evidence suggest that higher-order network interactions may be instrumental for spike propagation. For example, excitatory synaptic connectivity shows a prevalence of local neuronal cliques and patterns, and propagating activity in vivo displays elevated clustering dominated by specific triplet motifs. In this study we demonstrate a mechanistic link between activity propagation and higher-order motifs at the level of individual neurons and across networks. We algorithmically build spiking neural network (SNN) models to mirror the topological and dynamical statistics of neocortex. Using a combination of graph theory, information theory, and probabilistic tools, we show that higher order coordination of synapses is necessary for sustaining activity. Coordination takes the form of cyclic transitions between specific triangle motifs. The results of our model are consistent with numerous experimental observations in neuroscience, and their generalizability to other weakly and sparsely connected networks is predicted.

## Introduction

Network connectivity shapes dynamics in many systems and on many scales, ranging from gene transcription networks to epidemic spreading [1]. In the brain, neocortical architecture supports myriad complex functions. Fundamentally, before any of these functions can occur, signals must travel within and between local circuits and cortical regions. Thus spiking activity propagation is the most basic function that arises from the structure of synaptic connectivity in the brain.

Given the fact that the vast majority of excitatory synapses are weak and connections are sparse and recurrent, achieving stable activity propagation is highly non-trivial [2–5]. Theoretical and experimental studies have characterized several architectural features that have the capacity to promote and shape propagating activity, such as a long-tailed weight distribution, excitatory clustering and the balance between incoming and outgoing connections [3, 6–9]. Additionally, dynamical properties of ongoing activity, such as a balance between excitation and inhibition [8] and correlated spiking [10], are shaped by connectivity and in turn impact activity propagation.

Experimental results suggest that pairwise correlations alone may be insufficient to explain network dynamics such as propagation. Higher-order patterns in both structure and activity have been reported to be intrinsic features of neocortex [11] and may be key to our understanding of neuronal networks. Excitatory synaptic connectivity displays a prevalence of specific triplet motifs [2, 3] and cliques of neurons [8]. Propagating activity in real neuronal networks exhibit elevated clustering [12–18] that is dominated by triplet motifs which at least in part improves synaptic integration by coordinating the presynaptic pool [19]. Moreover triplet correlations are necessary to recapitulate spatiotemporal spiking patterns [20]. Computationally, they may improve coding [21, 22] and enhance perceptual accuracy and the prediction of responses in visual cortex [18, 23].

Spiking neural network (SNN) models are a promising avenue for studying the relationship between structure and function in neocortex. Here we use a novel algorithmic approach to build large numbers of sparsely-connected recurrent spiking neural network models to explore the role of higher-order interactions in activity propagation. These models are recurrent and sparsely-connected; they are comprised of excitatory and inhibitory adaptive exponential leaky integrate-and-fire (AdEx) neurons with conductance-based synapses [24]. Connectivity amongst excitatory units is clustered [8]. Network topology parameters are varied and informed by connectivity seen in cortex [3, 6, 7]. We performed grid search to tune network topological parameters to regimes characterized by asynchronous, low rate, and critical activity. Consequently our models closely approximate both the statistics of connectivity as well as spiking activity in neocortex [25–27]. We find that spiking during simulations of the same SNN topology can either spontaneously stop (truncate) or show sustained spike propagation (complete) on different runs, corresponding to different sets of initial membrane voltages, despite the fact that the runs exhibit epochs during which spike rates and critically are substantially similar. The dichotomy between sustained and truncated trials on the same networks provided us the opportunity to study the necessary conditions of sustained activity in spiking neuronal networks when they are not readily explained by spike rates, criticality or synchrony [28].

Using graph theoretic and probabilistic methods, we find that sustained activity requires cyclic higher-order coordination amongst excitatory neurons. In particular, the network cycles though epochs dominated in turn by three types of triangle motifs with low variance: fan-in triangle motif, followed by middleman and finally fan-out triangles [19]. When a network simulation fails to engage this low variance dynamical regime, spiking activity is not sustained and the simulation truncates. We find that strengthening weights and randomizing topologies in our networks lead to decreased clustering of units into triangle motifs. However, the relative motif ratios and transitions through time are maintained on sustained runs. Together, our results provide a mechanistic account and a possible explanation for the widespread findings of both clustered activity and synaptic connectivity in local neuronal circuits. The predictions of our model are consistent with numerous experimental measures in neuroscience and may be generalizable to other weakly interconnected networks that are not biological brains.

## Results

### Network construction and simulation

Each spiking neuronal network was comprised of 4000 excitatory and 1000 inhibitory adaptive exponential leaky integrate-and-fire (AdEx) units [24]. Synaptic connections were recurrent, sparse and conductance-based (Fig 1A). Excitatory connection strengths followed a long-tailed, log-normal distribution, where *µ* = −5.0 *\**10^−5.0^ nS, *σ* = .5 nS, corresponding to a mean of 1.13 nS and a variance of 0.365 nS (Fig 1B). Networks therefore had a large number of weak connections and few strong excitatory synapses. Connectivity in cortex is clustered [2, 16, 17]. Excitatory units were heterogeneously clustered, meaning that the number of units in a cluster varied (mean = 158.40, std = 12.27 units per cluster) [8]. The excitatory subgraphs had an average connection density of 0.211 (std: 1.10 *\** 10^−4^), of which 22.4% were recurrent (std: 0.02%). Total in-cluster density was 0.389 (std: 3.11 *\** 10^−3^) and out-cluster density was 0.196 (std: 9.94 *\** 10^−5^) (Fig 1C, D).

**Fig 1.**
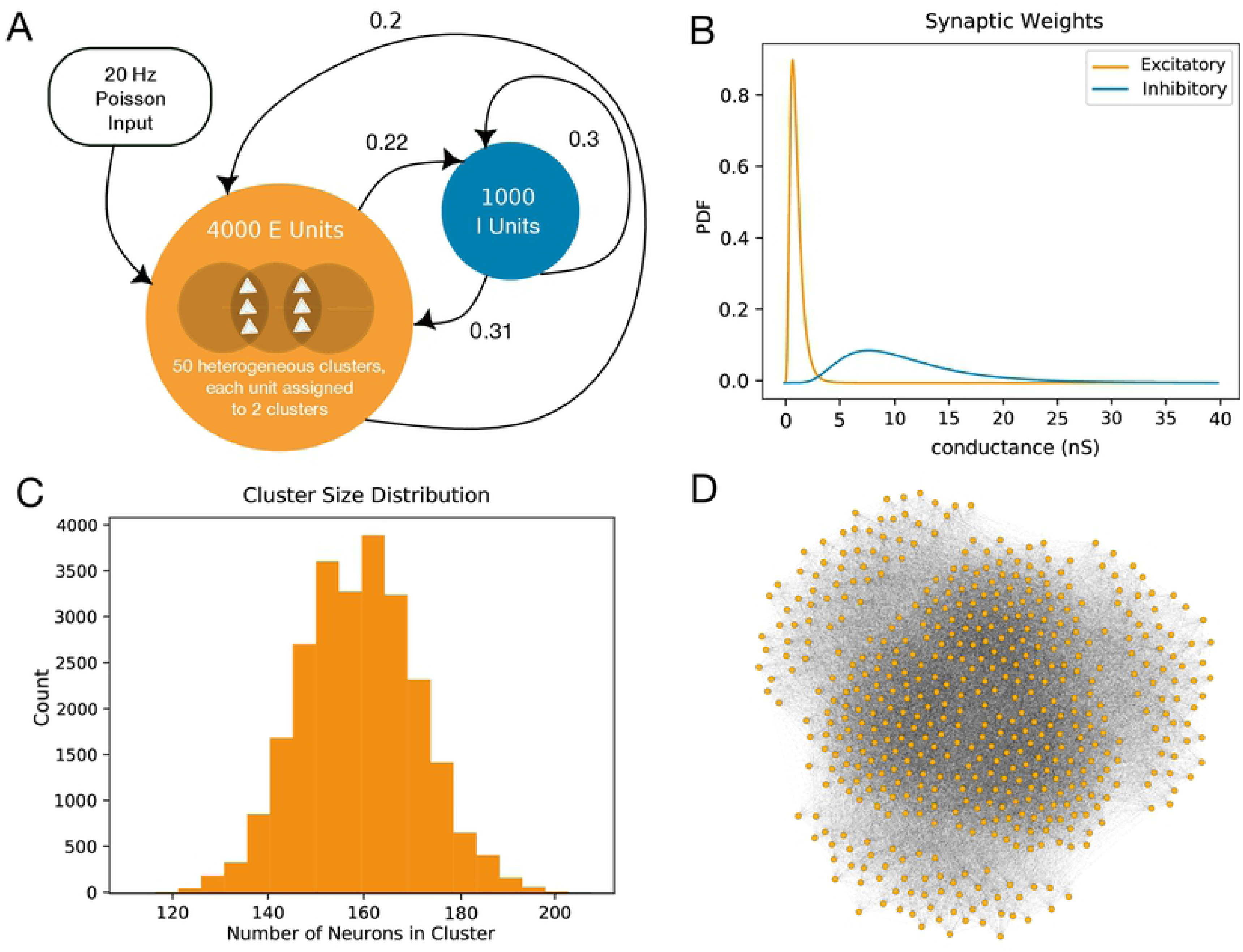
Network Construction and Search. A: Our networks were constructed with 4000 clustered excitatory and 1000 unclustered inhibitory units. Probabilities of connection were drawn from the literature and determined via grid search. Simulation runs began with 30ms of 20Hz Poisson input onto a subset of 500 units. B: Synaptic weights followed a log-normal (long-tailed) distribution. Synapses were conductance-based, so weights are in units of nanosiemens. Connections originating from inhibitory units were 10x stronger than those from excitatory units. C: For each network, we defined 50 clusters in total and randomly assigned each excitatory unit to two of these clusters. This resulted in heterogeneously-sized clusters. Here we show the cluster size distribution (in counts) for 500 networks. D: Visualization of a subset of 300 clustered excitatory units in our network.

At the beginning of a simulation trial, or run, initial resting membrane voltages were randomly assigned from a uniform distribution of -60 to -50 mV across all units. Activity was then initiated by 30 ms of 20 Hz Poisson input onto a set of 500 randomly chosen excitatory units (Fig 1A).

### Algorithmically identifying networks for analysis

In order to evaluate large numbers of networks while minimizing sampling bias, models were constructed, simulated, and scored algorithmically. We restricted the search for viable topologies to a range of connection likelihoods bounded by experimental observations [19]. This should not be interpreted to suggest that these connection likelihoods are the only viable solution for realistic spiking activity - we did not comprehensively survey the range of possibilities here.

We identified viable topologies iteratively; in the first iteration, we performed a low resolution grid search (Fig 2A). Specifically, we rewired topologies within a limited range of probabilities of connection from excitatory to inhibitory units,*p*_*e*→*i*_, and the probabilities of connection from inhibitory to excitatory units, *p*_*i*→*e*_, such that we identified sets of network connection likelihoods that resulted in topologies that successfully propagated spiking activity with low average firing rates and near-critical and low synchrony dynamics, as observed in neocortex [25–27, 29–32]. Criticality was measured using a branching parameter that is the ratio of active descendant units to active ancestor units across time [33]. A value of 1 - where the number of active descendants is equal to the number of active ancestors - indicates critical dynamics (Fig 2B). We used a fast, on-line synchrony heuristic (variance of the mean voltage divided by the mean of voltage variances, see Methods) for the sake of grid search speed. A run was considered to be asynchronous if this heuristic value was below 0.5. Runs below this threshold are shown to correspond to a high mean Van Rossum distance, a common measure of spike synchrony [34, 35](see Methods).

**Fig 2.**
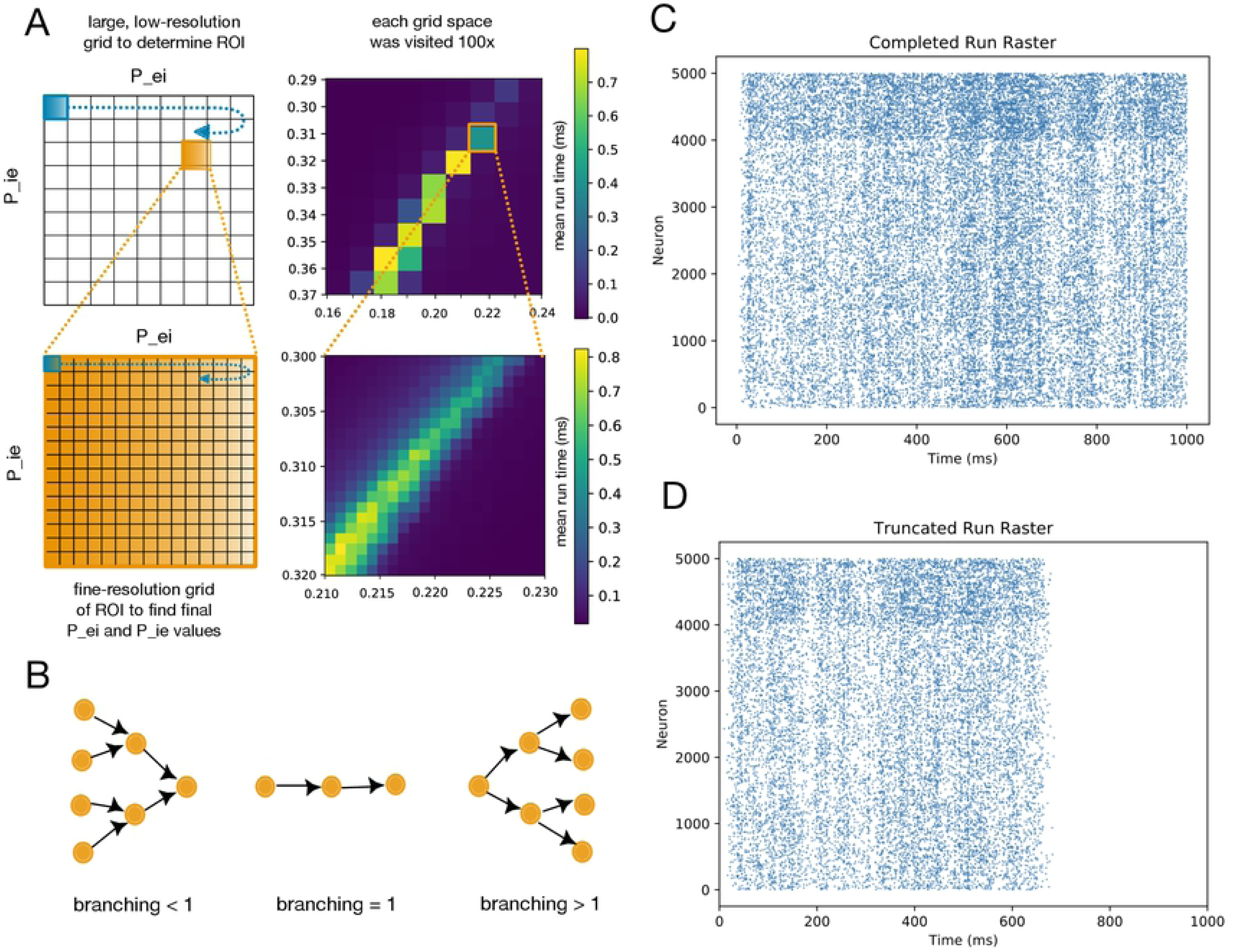
Comparison Between Scores on Complete and Truncated Runs. A: We performed two rounds of grid search for the topological parameters that yielded consistent low-rate, critical, and asynchronous dynamics. The first search was at a lower resolution to narrow down our region of interest, and the second was at a finer resolution. B: One scoring metric we used was branching. The branching parameter [33] is a proxy for criticality. It measures the ratio of active descendant units to active ancestor units. A branching value of 1 indicates a balanced (or critical) network, which is the value we optimized for. C: A raster plot of a single complete 1000ms simulation on one of our networks. Excitatory units are numbered 1-4000 on the y axis, and inhibitory units are 4001-5000. D: A raster plot of a single truncated simulation (700ms) on the same network with the same input.

The first iteration of grid search isolated a region of interest, and we next used a higher resolution grid to find specific topologies each with the same probabilities of connection but differing in the specifics of connections (Fig 2A). To find these topologies we used the best results obtained from the second round of grid search, which were *p*_*e*→*i*_ = 0.22 and *p*_*i*→*e*_ = 0.31. The values for *p*_*e*→*e*_ and *p*_*i*→*i*_ were taken from experimentally measured wiring probabilities in neocortex, and were 0.20 and 0.30 respectively [3, 19].

These connectivity parameters were used to generate 2,761 synaptic topologies, where each unique topology is referred to as a network. For each network we created 100 sets of input units, with 500 excitatory units per set. We ran 50 simulations on each set of input units, where each simulation starts with different membrane voltages for all units. Each simulation lasted for as long as spiking activity was sustained, up to a maximum of 1 second. The spiking activity of each run on each network was then scored according to the average firing rate, the level of synchrony, how balanced – or critical - the network was, and the duration of time over which spiking activity was maintained (see Methods). If a network’s average excitatory rate for all of its complete runs was less than 8 spikes/second, we added this network to the set of low-rate networks for subsequent analyses. High-rate networks were eliminated. This yielded a final count of 87 low-rate networks. For each of these networks, we determined the set of input units which led to the most consistently sustained simulations, with the trade-off of rate increasing slightly. We will refer to these as a network’s optimal input units. Optimal input units were used in generating 100 additional runs on each synaptic network, in which only the initial network state (i.e. membrane voltages of all units) varied. This generated a total of 8,700 unique runs, which we then analyzed.

### Scores on sustained and truncated simulations

We found that the same topology was capable of producing both sustained and truncated activity when only initial membrane voltages were varied. A run was sustained (or complete) if it displayed stable activity for the duration of a 1-second trial (Fig 2C). We found that all network simulations which reached 1 second were also able to sustain activity up to 10 seconds. We therefore chose one second as an indication of a network’s ability to sustain activity indefinitely, and as the definition of a successful run. If a network ceased all spiking before reaching the 1-second mark, that simulation was considered truncated or unsuccessful (Fig 2D). Scoring analysis of the network spiking dynamics of rate and branching between the two run types revealed significant overlaps.

We grouped truncated runs by their duration. Since network activity tended towards fewer spikes as a run approached truncation, we did not include the final 50ms of any run in the calculation of scores. We also did not consider the stimulus period (initial 30ms), as we wished to analyze self-sustained network dynamics rather than stimulus-driven spikes. By focusing our analyses on the middle portion of each run, we find that the rate and branching values of both sustained and truncated populations overlapped substantially. Runs that truncated at 100ms, at 500ms, and runs that were sustained for greater than 990ms shared similar mean excitatory firing rates (9.95, 9.77, and 10.14 spikes/s, respectively). Runs that truncated between 140 and 400 ms tended to have a higher mean rate (15.65 spikes/s), demonstrating that higher firing rates can contribute to instability of a network [28]. The overlap index between truncated run rate and completed run rate was measured to be .27. The overlap index for criticality for the two run types was measured to be .31. In contrast, the synchrony measure was much more discerning between the run types, yielding an overlap index of .0021. We found that Van Rossum spike distance increased (synchrony decreased) as run duration increased. This inverse relationship between synchrony and run duration goes against the intuition that successful spike propagation is at least in part due to synchronous activity [36–42]. Regardless, it was clear that first order spiking statistics did not provide simple explanations of how and why activity was sustained in some cases and truncated in others.

### Graph theory analysis of simulated networks

We considered functional interactions between neurons to provide an explanation for sustained activity. To do so we analytically evaluated the networks using graph theory. In previous work we have defined a taxonomy of active networks to focus our analysis [19]. We refer to the structural connectivity matrix of our models as the synaptic graph (Fig 3A). From each simulation on a synaptic graph, we generated a single functional graph as well as a time series of recruitment graphs at 10ms resolution.

**Fig 3.**
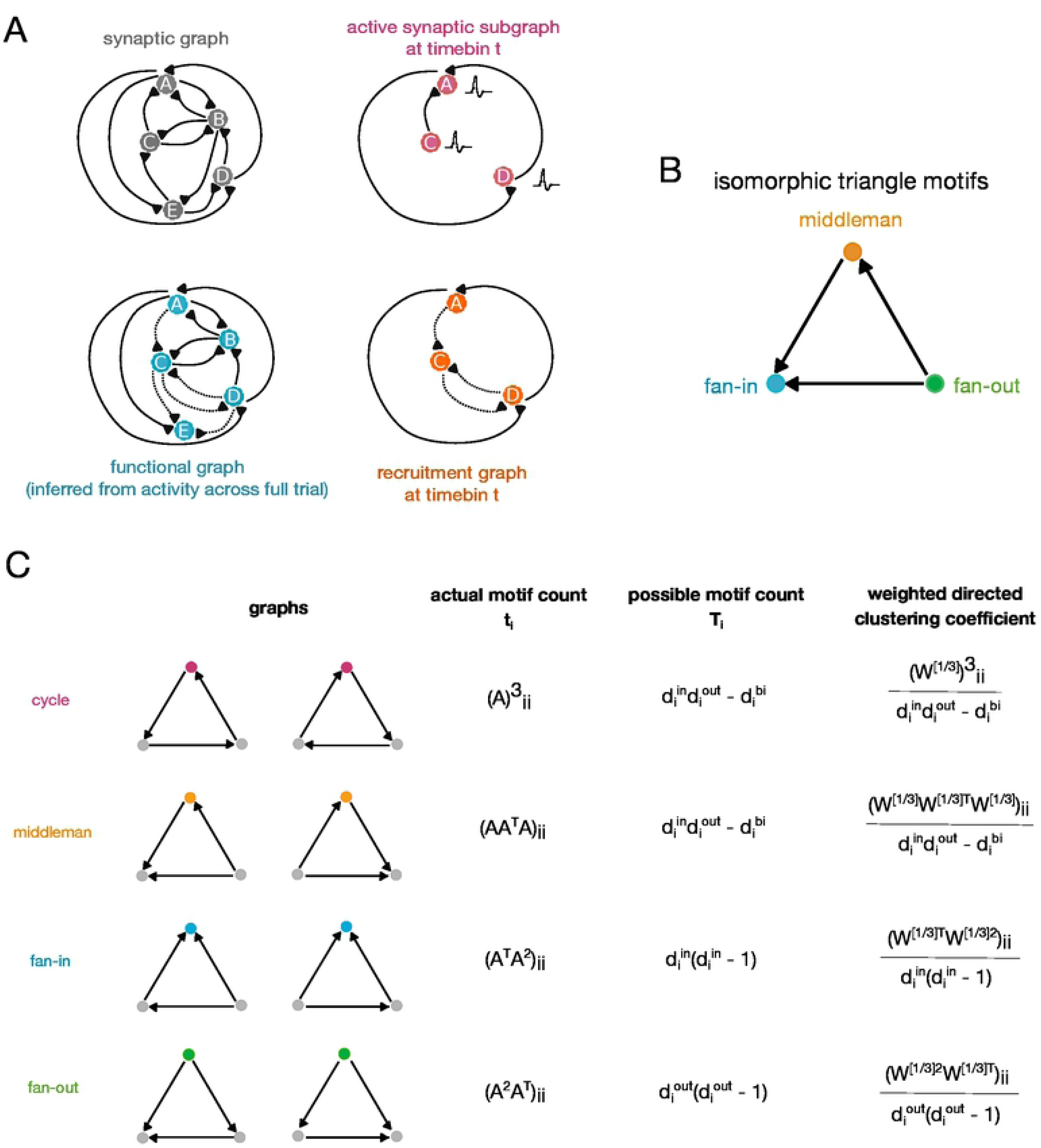
Graph and Motif Definitions. A: The synaptic graph is the ground-truth topology of our networks. Based on spiking activity during each simulation, we construct a series of active synaptic subgraphs - one for each time bin. These are graphs made of units which spiked in that bin, connected via the same edges as in the synaptic graph. We infer a single functional graph from whole-trial spiking activity using confluent mutual information - these graphs represent the functional connectivity of the network for each simulation trial. The intersection of the functional graph with the active subgraph for a given time bin yields the recruitment graph for that time bin. B: The three triangle motifs we examine - fan-in, fan-out, and middleman - are isomorphic by rotation. When calculating motif clustering, the choice of reference node is key. C: Calculation of the clustering coefficients of the different triangle motifs on weighted directed graphs, as defined in Fagiolo 2007. The clustering coefficient is defined as the ratio of the actual to the possible motif counts.

We constructed functional graphs using mutual information to quantify pairwise correlations between spiking neurons across each simulation. In order to generate a series of recruitment graphs we identified the intersection of the functional graph with the synaptic subgraph according to the units which were active in each 10ms time step, resulting in one recruitment graph per time step. Weight values of functional and recruitment connections were calculated from mutual information and summarized in the functional graph, rather than taken from the synaptic weight matrix (see Methods). Because of our interest in the relationship between synaptic structure and functional spike propagation, we focused our analysis on recruitment graphs.

### Triplet Motifs

The term ‘motif’ refers to a pattern formed by a group of units in a network. Previously we found that triplet motifs were informative of synaptic integration [19] and also increased the power of encoding models [18, 21–23]. Here we focused our analysis on similar patterns of connectivity in the recruitment network, involving groups of three units [43].

From the perspective of a single reference neuron, neighboring neurons can be arranged into four types of triplet motifs: fan-in, fan-out, middleman, and cycle. In isolating one triplet, the fan-in, fan-out, and middleman motifs are isomorphic by rotation, meaning that they only differ due to the choice of reference node (Fig 3B). The relative importance of a motif for a given neuron is measured by its contribution to that neuron’s clustering coefficient (Fig 3C). The clustering coefficient is the weighted ratio of the actual over the possible counts of a particular triplet motif type in which that neuron participates. Individual reference nodes in a given triplet may yield different clustering coefficients due to their specific weights and connections (see Methods).

It was possible that each of the algorithmically generated networks had different connection densities and weight distributions, which would impact weighted motif clustering coefficient measures. In this case a measure that incorporated weight and controlled for density would be especially relevant since the recruitment graph density changes in time. We therefore used the measure of clustering propensity [44]. Propensity is the ratio of the clustering coefficients of the recruitment graphs compared to the average clustering coefficients of graphs with the same connection structure but randomly assigned connection weights. The propensity measure allowed us to compare different networks despite different connection densities and also allowed us to assess the impact of specific edge weights on triplet motif clustering coefficients [19, 44]. A propensity value of 1 indicates that specific edge weights play a negligible role in clustering, since random edge weights would yield the same clustering coefficients (see Methods).

### Density and recurrence statistics

As reported above, synaptic networks were 21.1% connected, and 22.4% of connections were reciprocal (Fig 4A, B). The functional networks of complete runs, which were calculated using mutual information and were unique to each run, were more dense and also more recurrent. The functional networks averaged 32.6% (std: 0.6%) connectivity, of which 59.0% (std: 0.4%) were recurrent. Recruitment graphs across time in complete runs were sparser than the synaptic graphs, although only slightly less recurrent (9.5% connected, std: 0.5%, and 16.7% recurrent, std: 0.5%). Functional and recruitment density and recurrence did not differ significantly between complete and truncated runs.

**Fig 4.**
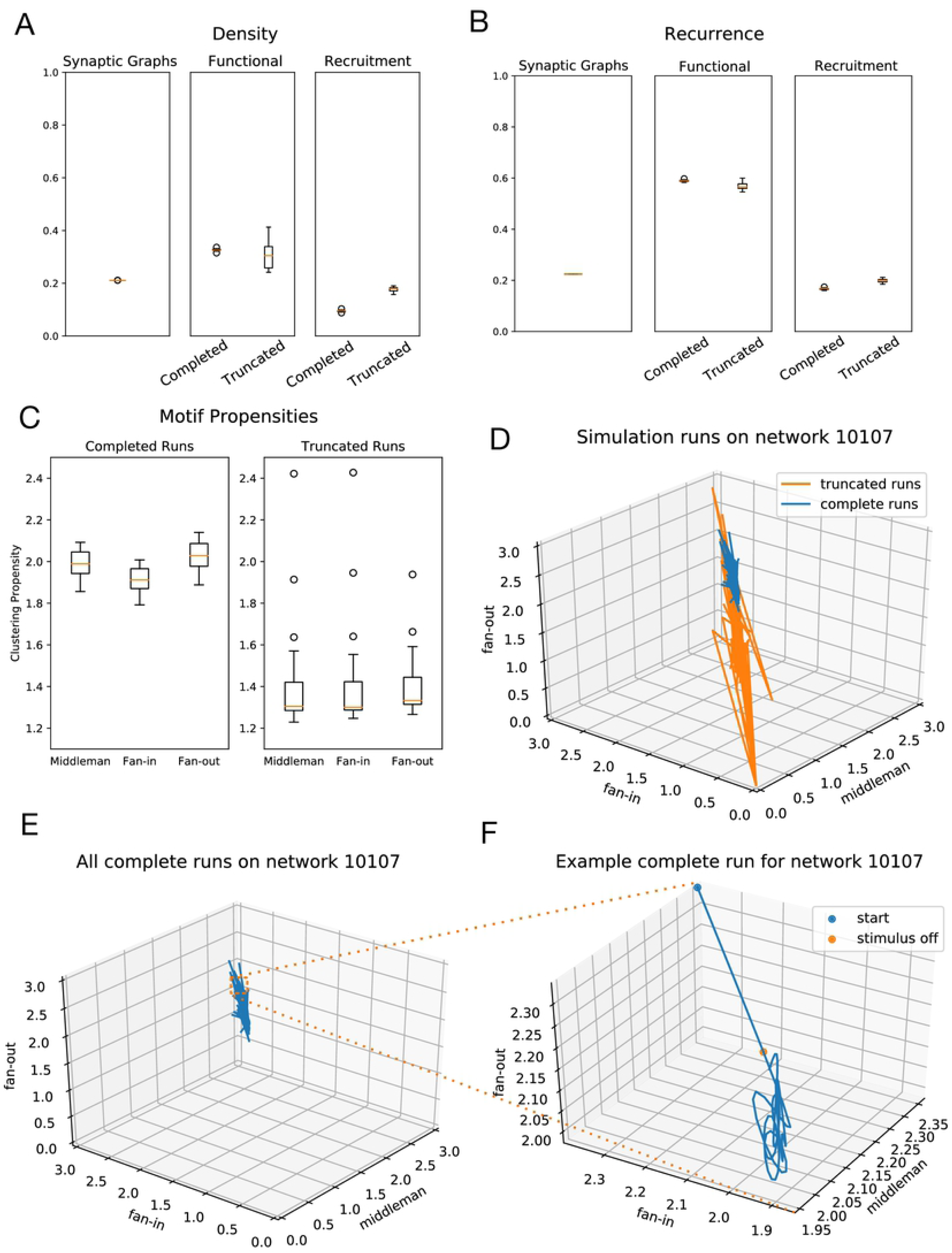
Standard Network Triplet Motifs. A: Density (ratio of existing to possible connections) of synaptic, functional (complete vs truncated), and recruitment (complete vs truncated) graphs across all networks. B: Proportion of existing connections which are recurrent in synaptic, functional (complete vs truncated), and recruitment (complete vs truncated) graphs across all networks. C: Comparison of isomorphic triangle motif clustering propensities on complete and truncated runs across all networks. D: Trajectories of all runs on a sample network in 3-dimensional isomorphic motif space. Truncated runs have a larger spread of trajectories and are shown in orange, complete runs are shown in blue. E: Trajectories of all complete runs alone, on axes of the identical scale as in panel d. F: Example trajectory of a single run on the same network, now enlarged (from inset in panel e). The network begins away from the area of its eventual cyclic trajectory, and the 30ms of Poisson input at the beginning of the run drives it towards this region.

### Triplet motifs in the different graph types

We found that the three isomorphic motifs showed equal clustering in the synaptic graphs. This is expected of graphs with random, albeit clustered, synaptic connectivity. Clustering propensity centered at 1.00 (std = 7.8 *\**10^−5^, 7.9 10^−5^, 8.0 *\**10^−5^ for middleman, fan-in, and fan-out) for all three isomorphic motifs (Fig 4C). A value of 1 indicates that specific edge weights in synaptic graphs play a negligible role in clustering, since random edge weights would yield the same clustering coefficients.

We found that in contrast to the static synaptic graph the dynamic functional and recruitment graphs were not random. The isomorphic motifs’ dominance in the recruitment graphs, or the strength of each motif’s contribution to overall clustering, varied over time in each trial. For complete runs, motif clustering propensities for recruitment graphs (averaged across all time and all topologies) were 1.98 (std = 0.06), 1.91 (std = 0.06), and 2.03 (std = 0.07) for middleman, fan-in, and fan-out, respectively. Propensity values greater than 1, as these are, indicate that units in the recruitment graphs are more strongly clustered than would be expected in structurally-matched graphs with randomized weights. Isomorphic motif clustering propensities also varied in recruitment graphs of incomplete runs, with averages of 1.39 (std = 0.23), 1.40 (std = 0.23), and 1.43 (std = 0.24) for middleman, fan-in, and fan-out motifs (Fig 4C).

### Cycling of triplet motifs

To evaluate how the three isomorphic motifs co-varied across time for both successful and truncated trials, we plotted motif clustering propensities at each point in time against one another (Fig 1D, E, F). Clustering propensities formed a cyclic trajectory within a restricted region of motif space. This indicates a systematic alternation between over- and under-representation of the three isomorphic motifs in the whole network relative to what would be expected in edge-matched networks. We found that the cyclic trajectory within this region of motif space was consistent for all low-rate, excitatory clustered networks we examined. We also found the same orderly temporal progression from one isomorphic motif to another when we considered clustering coefficient values. In contrast, truncated simulations were never restricted to this behavior.

### Motif cycling and sustained activity

The motif cycling trajectory was not present at the moment of first spikes in a simulation. Rather, the path started at a point in motif space as determined by the initial membrane voltages of all neurons in the network. Injection of Poisson input drove network activity towards its eventual trajectory (Fig 4F). We identified two distinct types of truncation - in the first and far more common (97.7%) of the two, the simulation trajectory never approached or entered the region in propensity motif space where sustained runs lay. In the second, rarer case, the simulation successfully entered the sustained regime, yet after several hundred ms the trajectory destabilized, resulting in truncated activity. Truncated trajectories did not follow a canonical path. Instead, motif dynamics during truncated runs transited in all directions away from the central region, demonstrating the multitude of ways in which activity structure can lose stability resulting in a failure of spike propagation (Fig 4D).

We examined whether the initial distance and input trajectory, which were determined by the initial conditions of the network and the Poisson stimulus, were determinants of successful activity propagation. We found that even if the distance from the cycling region at the end of the stimulus period was minimal, some simulations still failed to enter into and stay within that regime. Others which were still distant from the region after the initial stimulus period continued on a successful trajectory and entered a stable cycling regime. These behaviors point to complex interactions between the network’s internal state and how input onto precise units within that network can influence propagation.

### Markov Analysis

In order to quantify the cycling between isomorphic motifs, we constructed a Markov model for state transitions between dominant isomorphic motifs. We described the network using a probabilistic voting scheme, as opposed to using analog propensity values. A unitary vote is cast by each unit for the motif type for which it has the highest propensity value at some time step. The proportion of total votes for each motif type is used to describe the relative dominance of that motif at that time step.

From this time series we constructed a Markov model transitioning between states. We found that the parameters characterizing the Markov process were canonical and low variance, such that successful cycling followed a specific reliable sequence between motifs. In contrast, the Markov parameters in simulations that truncated showed a failure to recruit this low-variance canonical sequence. First and second-order state probabilities and state transition probabilities significantly differed between completed and truncated runs (p *<* 0.001) (Fig 5A, B). Second-order state conditional probabilities also differed (p *<* 0.05) (Fig 5C). State probability is the probability of a motif being dominant at a given time. Second order probability is the probability with which a sequence of two motifs will be dominant at some given time. Conditional probability is defined as the probability of a motif given history of previous two motifs.

**Fig 5.**
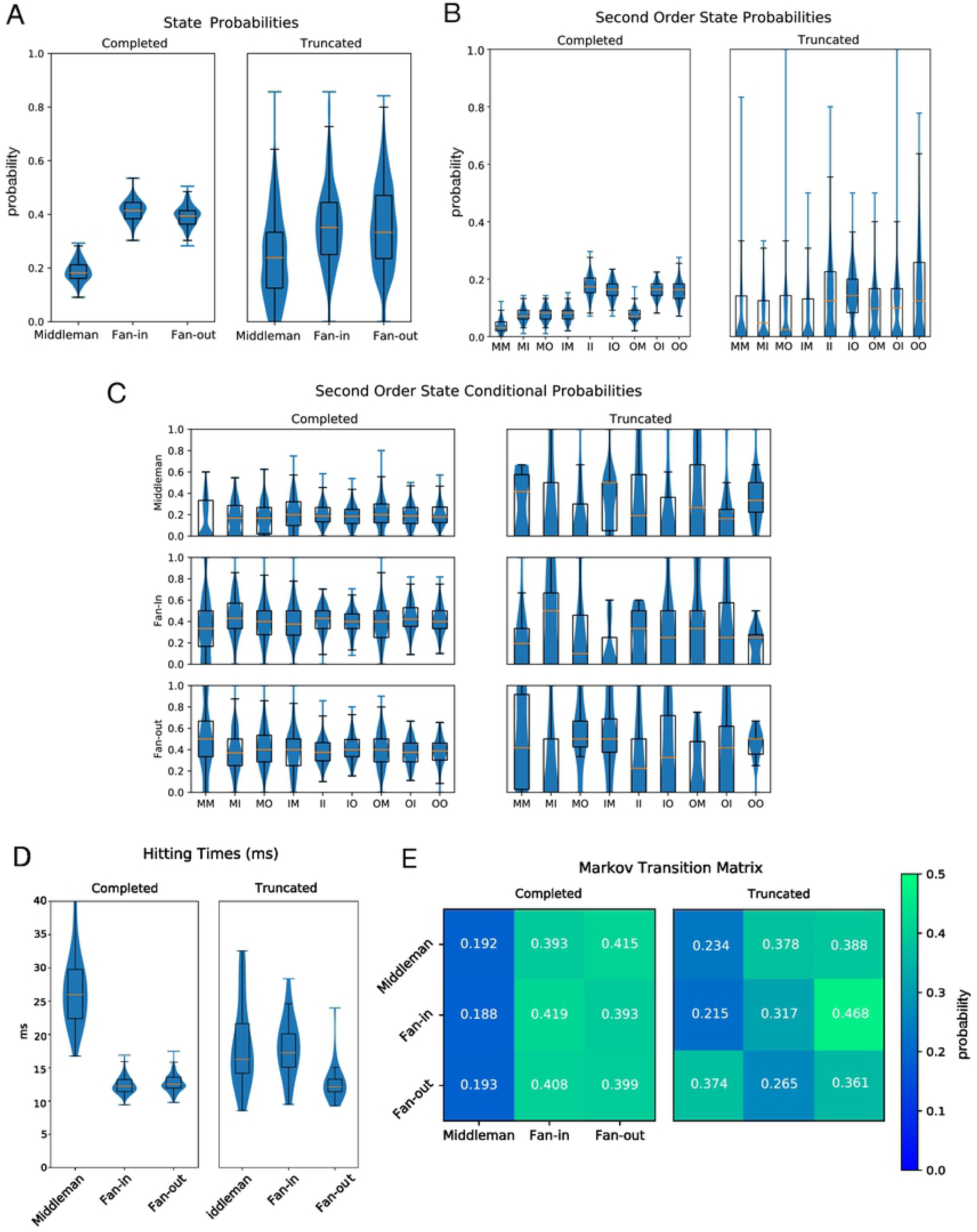
Markov Comparisons Between Completed and Truncated Runs on Standard Networks. A: Probabilities of state dominance of a triplet motif in completed (left) and truncated (right) runs. B: Second order state probabilities for completed (left) and truncated (right) runs. C: Second order conditional state probabilities for completed (left) and truncated (right) runs. D: Expectation of hitting time for Markov model of state dominance transitions in completed (left) and truncated (right) runs. E: Visualization of Markov matrix for state dominance in complete (left) and truncated (right) runs.

Markov analysis also gave the time scale which characterized motif cycling via the mean time for recurrence. This is defined by the expectation of the hitting time for each motif, given the network is currently dominated by that motif. We define hitting time, *t*, as *H*_*i*_ = inf *{n ≥* 1 : *S*_*n*_ = *i*|*S*_*o*_ = *i}* and our expectation of hitting time, *t*, as 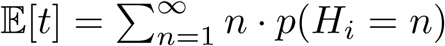. This gives a mean recurrence time for each motif. We find truncating middleman to have mean 18.59 ms (std: 6.49 ms), completing middleman to have mean 27.02 ms (std 5.99 ms), truncating fan-in to have mean 18.01 ms (std: 4.61 ms) completing fan-in to have mean 12.39 ms (std: 1.27 ms), truncating fan-out to have mean 12.95 ms (std: 3.14 ms), and completing fan-out to have mean: 12.65 ms (std: 3.56 ms). Hitting times differed significantly between completed and the small subset of truncated runs that entered this region of propensity (p *<* 0.001) (Fig 5D).

### Effects of connectivity weights

We hypothesized that the cycling between clustering propensities was necessary for sustained activity due to the weak strength of the majority of individual synapses. Fan-in clustering has the highest probability of remaining in the state of fan-in clustering in the next time point which hints at the greater need for integration. But once integration is sufficient, the motif changes. For our model and for most of the synapses in neocortex, convergence of spikes from multiple sources must occur in order to evoke spikes in a receiving neuron [19]. Consequently we expected that as connection weights increased, the cycling between population-wide isomorphic motifs would lessen.

To test this, we strengthened all synaptic weights in the networks that previously scored well from 1.0x to 2.0x original values in increments of 0.1. Simulations were then re-run on these strengthened networks using the same stimulus and initial conditions. At 1.6 times the original weights, networks consistently displayed bursting activity. Consequently we restricted our analysis to networks with weights increased 1.5 times. All runs on these networks reached completion.

Increasing weights led to a decrease in all triangle motif propensities, and also led to differences in the Markov characterization. Motif state probabilities differed significantly between completed runs on the original graphs and those on graphs with increased weights (Chi-square test, p *<* 0.001). Second-order state probabilities, state conditional probabilities, hitting times, and state transition probabilities all differed significantly as well (p *<* 0.01, p *<* 0.05, p *<* 0.005, p *<* 0.001 respectively), demonstrating the interaction of synaptic reliability on the necessity of this regime (Fig 6). However the trend remained and in all sustained runs a low variance transition from motif to motif occurred.

**Fig 6.**
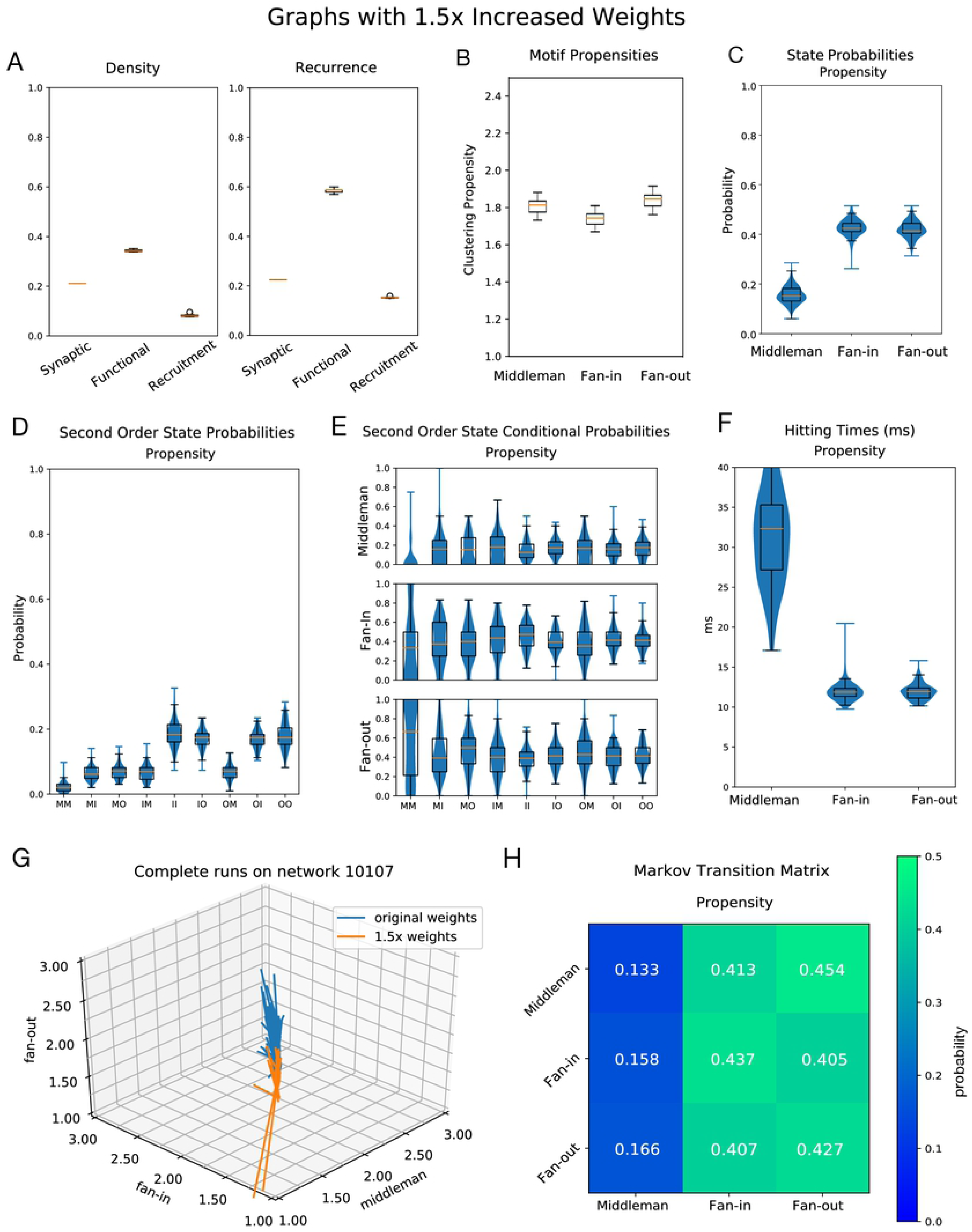
Networks with Increased Weights. Networks have the same structure as those seen in figures 4 and 5, but all edge weights have been increased by 1.5 times their original values.A: Left, density (ratio of existing to possible connections) for synaptic, functional, and recruitment graphs. Right, recurrence (ratio of recurrent to all existing connections) for synaptic, functional, and recruitment graphs. B: Clustering propensity for isomorphic triangle motifs on increased-weight-graph simulations. The y-axis is scaled to match that of Figure 4C (clustering propensities on original graphs) and Figure 7B (clustering propensities on unclustered ER graphs). C: Probabilities of dominance of each triangle motif. The dominant motif at a time point is given by the maximum of mean middleman, mean fan-in, and mean fan-out across units. D: Second order motif state probabilities for progression of temporal recruitment graphs. E: Probabilities for each motif to follow a given second order motif. F: Hitting times for each state for the Markov process defined by motif transition probabilities. G: Trajectories of all complete runs on a sample network in 3-dimensional isomorphic motif space. In blue are the runs on the network with its original weights, in orange are the runs on the same network with weights increased. H: Markov Matrix for transition probabilities between motifs.

### The dynamical motif solution is arrived at regardless of synaptic connectivity statistics

The networks on which we performed all our analyses have excitatory clusters of units. To test whether our results, including the motif cycling phenomenon, are dependent on this structure, we next examined non-clustered Erdős-Renyi (ER) graphs with *p*_*i*→*e*_ = .25 and *p*_*e*→*i*_ = .35. ER graphs had the same *p*_*e*→*e*_ and *p*_*i*→*i*_ values as the clustered networks. We found that transitions between motif types were also present in the activity of sustained runs on ER networks (Fig 7). The relative increase in clustering in ER graphs when comparing synaptic to recruitment graphs is substantially greater than seen in our graphs with excitatory synaptic clusters. In the synaptic networks, triplet clustering coefficients average 0.11. However, this value increased to 0.20, 0.09, and 0.15 for fan-in, fan-out, and middleman motifs in the recruitment graphs. The propensity values for all isomorphic motifs were consistently lower than 1, with means centered at 0.8 (Fig 7). We find that unclustered ER graphs and clustered ER graphs differ significantly (p*<*.005) in second-order state probabilities, state conditional probabilities, hitting times, and state transition probabilities. As in the case with the increased weights however the qualitative cycling of motifs was present in sustained runs.

**Fig 7.**
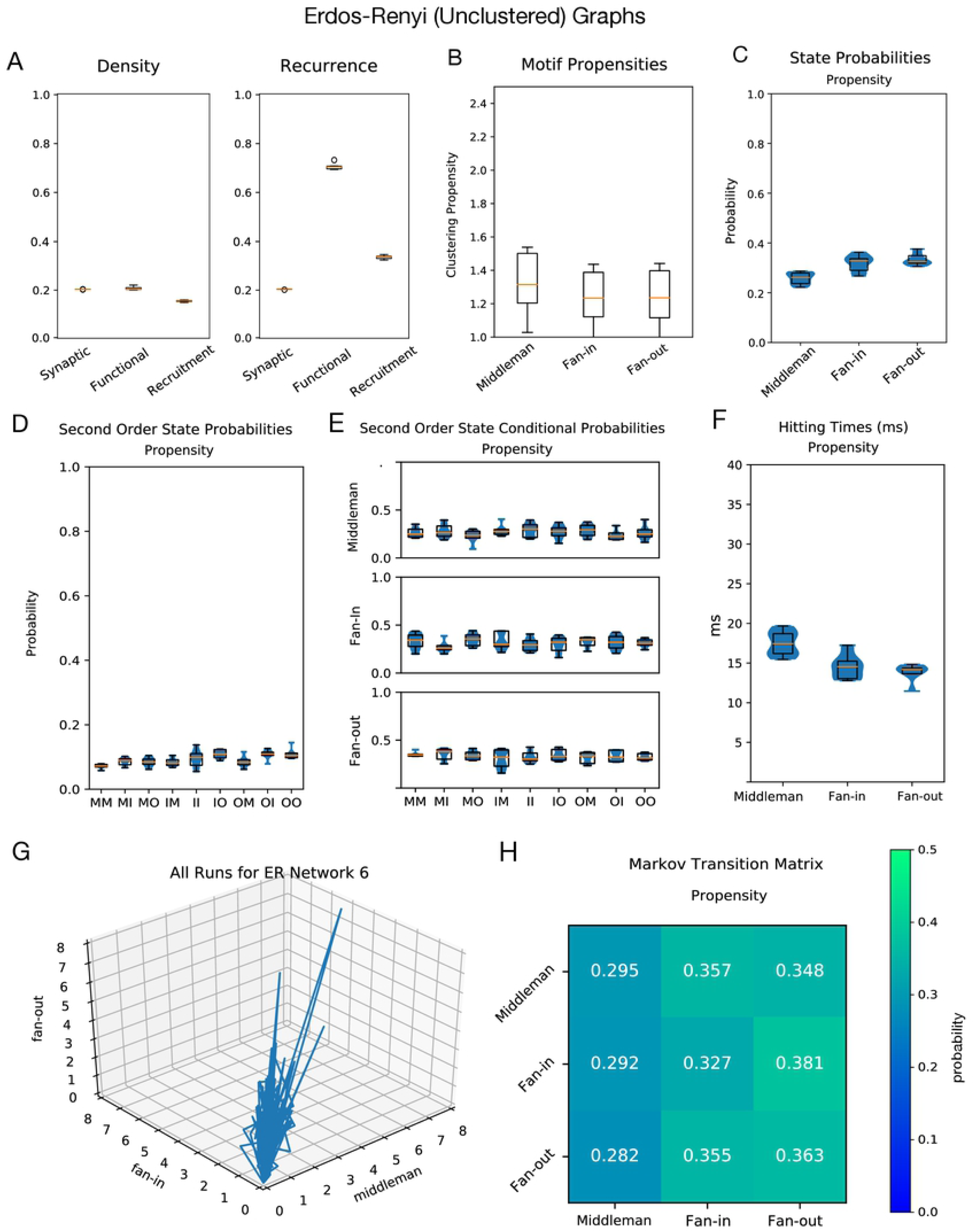
Unclustered (Erdős-Renyi) Networks. A: Left, density (ratio of existing to possible connections) for synaptic, functional, and recruitment ER graphs. Right, recurrence (ratio of recurrent to all existing connections) for synaptic, functional, and recruitment ER graphs. B: Clustering propensity for isomorphic triangle motifs on ER graph simulations. The y-axis is scaled to match that of Figure 4C (clustering propensities on original graphs) and Figure 6B (clustering propensities on graphs with 1.5 times increased weights). C: Probabilities of dominance of each triangle motif. The dominant motif at a time point is given by the maximum of mean middleman, mean fan-in, and mean fan-out across units. D: Second order motif state probabilities for progression of temporal recruitment graphs. E: Probabilities for each motif to follow a given second order motif. F: Hitting times for each state for the Markov process defined by motif transition probabilities. G: Trajectories of all runs on a sample ER network in 3-dimensional isomorphic motif space. All runs reached completion. H: Markov Matrix for transition probabilities between motifs.

## Discussion

This work demonstrates that higher-order structure is required for sustained activity in low-rate recurrent networks such as neocortex. Spikes must traverse the network in a coordinated way, cycling between the dominance of three triplet motifs. The transitions between fan-in, middleman, and fan-out motifs reveal the necessity of balance between distribution of signal and convergence of inputs necessary to integrate those inputs. The presence of these motifs in the recruitment graphs reflects the functional routing of activity through synaptic connections. When synapses become stronger and more reliable, overall triplet clustering decreases, while the reliability of their transitions remains. Higher order motifs in the recruitment network thus provide a direct link between network activity and the stable activity in that network.

Simpler measures, such as rate or synchrony, did not easily explain network stability in our models. Previous work has demonstrated that sustained, asynchronous network activity co-occurs within a specific range of firing rates, supported by a balance between excitatory and inhibitory conductance [28]. Counter-intuitively, networks were unable to continue to spike both when firing rates are low and also when rates are too high. We also found a counter intuitive result in that networks which display asynchronous spiking dynamics tend to sustain activity for longer than networks with more synchronous spikes. Previous work suggests that synchronous spikes are necessary and particularly efficacious for signal propagation to occur [36–42]. For example, one proposed mechanism for communication between two regions is via synchronous spike volleys with coherent phases, which capitalize on the recurrent nature of connections in neocortex [38]. Yet synchrony is intricately tied with rate - transmission speeds are higher when spikes are more synchronous. High-rate and high-synchrony spikes may overwhelm the integrative capacity of downstream units, contributing to instability. Our results lend nuance to a previously-held view that although neocortex displays asynchronous activity, it hinders rather than helps spike transmission. We suggest that certain levels of asynchrony are in fact necessary for spike transmission at the network level just as in the case with firing rate.

In addition to being low-rate and asynchronous, our networks were algorithmically constructed to have critical dynamics. A system can be described as critical if it operates near a phase transition or “critical” point. A classic example is a mound of sand, which is said to be at the critical point if each new grain of sand added causes an avalanche of sand following a power law distribution. In neocortex this entails activity that, in the absence of external input, propagates activity without become epileptic or dying off, and which follows a power law distribution in its active population size [33]. However, these are only necessary conditions and not deterministic of all critical systems. The idea that neocortex operates near a critical point has a long history in neuroscience, going back to Alan Turing [45], and has been implicated in a number of desirable properties for neural networks [46]. For example, networks tuned near the critical point display maximum information transmission [33], information storage [47], and computational power [48]. Understood as a branching process, critical activity entails very little structure in activity, namely that the average ratio of descendants to ancestors be 1.

Our results provide an explanation for the prominence of higher order motifs in real data. Elevated motif counts have been observed in synaptic connectivity and in recordings of clustered activity in vivo [2, 3, 8, 12–18]. Through mechanisms of learning in neocortex such as STDP, functional patterns may be further strengthened to enhance integration in cortex. We wish to draw attention to the fact that our study focused on the whole-network scale. Individual units spiked only sparsely, making it difficult to continuously track single-unit motifs across small epochs of time since interspike intervals were generally larger. The models we used were constructed to simulate neocortex. The network structures we employed closely match experimental observations [2, 3] and the model units capture many of the statistics of neocortical neurons [5]. Our results provide, first and foremost, an account of the role of beyond-pairwise interactions in the brain. Yet the behavior of these models may reflect necessary features of weakly-connected networks in which integration from multiple sources is necessary for the system to succeed. In such systems it is likely that stability relies on higher-order patterns. For example, the spread of rumours in a social network relies on integrating interactions. Social networks are small-world networks characterized by clusters, a feature which is present in our model as well as many other systems [49, 50]. The “illusion-of-truth” effect in rumour spreading on a social network has the integrate-and-fire property, where an individual may need to hear a rumour from multiple sources before they reach a confidence threshold to repeat it to others [51].

The necessity of higher-order patterns for stable activity has strong implications for neural coding. Previous work has already demonstrated that correlations enhance coding, with triplet correlations having an advantage over pairwise [5, 16, 18, 21–23, 52–54]. The neural code must rest upon a foundation of stable propagation of spikes, which we have shown in turn rests on higher-order motifs and coordinated integration. Any two spikes must take place within some time interval for them to interact. The asynchronous and critical regime observed in vivo and in our models pushes the limits on what constitutes a cooperative event. In our model, the precise conditions for integration are dictated by the time constants we chose, while in neocortex the same time constants may vary and span some range. Neuromodulation, cognitive state, and a variety of other factors all dictate the requirements which need to be met for integration. Local connectivity certainly plays a large role as well. Consequently the role of higher order interactions in coding and in coordinating synaptic integration may vary by state.

## Materials and methods

### Network Structure

Our graphs are recurrent and sparsely connected networks of several thousand adaptive exponential leaky integrate-and-fire (AdEx) units with an extra poisson input term [24]. Synapses between all units are conductance-based. This enhances realism by taking neuron-specific state features into account during synaptic integration [24]. Specifically we define our neuron Voltage, *V*, as.

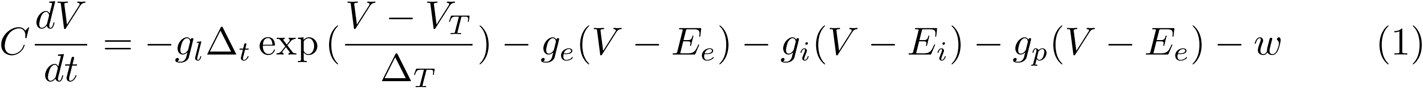

adaptation current, *w*, as

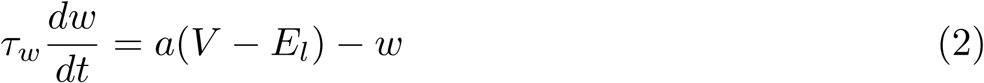

excitatory conductance, *g*_*e*_, as

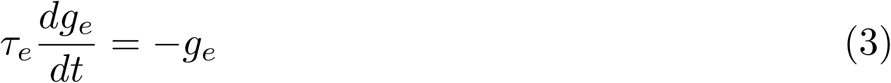

inhibtory conductance, *g*_*i*_, as

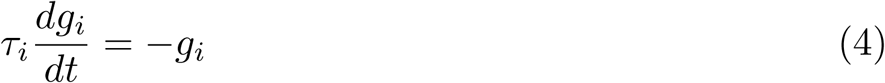

poisson input conductance, *g*_*p*_, as

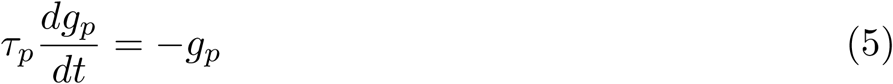

Spike was said to occur if *V > V*_*t*_, at a spike *V* was set to *EL, w* was incremented by *b* and *g*_*e*_ and *g*_*i*_ were incremented by synapse weight if downstream of the spiking neuron.

For information on parameters, see S1 Table. Each network is comprised of 1000 inhibitory and 4000 excitatory units. Precise wiring probabilities between excitatory and inhibitory populations were determined through grid search within biological constraints.

Network synaptic connectivity is heterogeneously clustered [8]. For each network we defined 50 total clusters, with each excitatory unit randomly assigned to two clusters. Clusters thus vary in size and follow a normal distribution. The wiring probability between two units within the same cluster is twice that of units in different clusters. Network cluster sizes range from 111 to 207 excitatory units (mean = 158.40, std = 12.27). Inhibitory units are not clustered; their wiring probability is uniform across the graph.

Edge weights follow a long-tailed distribution (Fig 1B). Edge weights that originate from inhibitory units have conductances which are ten times greater than those which originate from excitatory units, in accordance with experimental results [55].

### Network Simulation

Each simulation was recorded at 0.1-ms temporal resolution. A trial began with 30 ms of Poisson input stimulus onto 500 randomly chosen units. After 30 ms the stimulus would cease and activity would propagate naturally through the network. The simulation would continue for as long as spiking activity is sustained, up to a maximum of 1 second. If during a simulation no spikes occur across the network for 100 ms, the network is deemed inactive and the simulation trial is halted. We found that all network simulations which reached 1 second were also able to sustain activity up to 10 seconds. We therefore chose one second as the marker for a network’s ability to sustain activity indefinitely, and as the definition of a successful run. Upon completion each simulation yields an output raster of spike times for every unit in the network. The poisson input train, input units, network topology, and initial conditions of all units were recorded for each simulation. This enabled subsequent analyses and also allowed for re-use of a synaptic graph or re-instantiation of a simulation using some of the original settings while varying others.

### Parameterization

Our models are constructed to parallel the features of biological neural networks, but are also constrained by considerations of computing resources. In a study which modeled cortex with high biophysical and anatomical detail, simplifying the neuron model did not lead to drastic differences in the network’s behavior from the detailed model or from in vivo results. Most qualities remained unchanged, suggesting that in many cases extremely granular models are not necessary to yield experimental insights [24]. Instead, the most important feature for retaining qualitative correspondence are the rules of synaptic connectivity. Therefore we required our models’ connectivity parameters to closely match those of biological neural networks.

The probabilities of wiring between excitatory (E) and inhibitory (I) populations in our models were taken directly from or bounded by the results of biological experiments. The wiring probabilities from E to other E units and from I to other I units in neocortex are well-studied, but there is less data on connections from E to I and from I to E. We therefore used an algorithmic approach to find the optimal values. Beginning within a biological range, we used grid search to find values of *p*_*e*→*i*_ and *p*_*i*→*e*_ that led to successful propagation of activity at the lowest possible rates. We used these optimal wiring rules to construct all synaptic graphs in this study.

Two iterations of grid search were used to find the wiring parameters needed to maintain naturalistic spiking for the duration of a simulation (Fig 2A). We searched for the optimal probability of connection from excitatory to inhibitory units, *p*_*e*→*i*_, and the optimal probability of connection from inhibitory to excitatory units, *p*_*i*→*e*_, such that networks would propagate activity at the lowest possible rates. In the first iteration, we used a low resolution grid (space size 0.001) to search for *p*_*e*→*i*_ within the range 0.16 to 0.24 and *p*_*i*→*e*_ within the range 0.29 to 0.37. These two ranges were taken from known wiring probabilities in neocortex. Each grid space was visited ten times to achieve an average measure of rate and completion. This isolated a region of interest where the rate was lowest, between *p*_*e*→*i*_ values of 0.216 and 0.220, and between *p*_*i*→*e*_ values of 0.309 and 0.313. We used a higher resolution grid (space size 0.0001) to explore this region.

For all subsequent simulations we used the best results obtained from grid search. The optimal probability of wiring for excitatory to inhibitory units, *p*_*e*→*i*_, was found to be 0.22, and the optimal value for *p*_*i*→*e*_ was 0.31. The values for *p*_*e*→*e*_ and *p*_*i*→*i*_ were taken from known wiring probabilities in neocortex, and were 0.20 and 0.30 respectively [19]. Based on these wiring rules, we constructed synaptic graphs of networks for simulations. Each synaptic graph is a matrix *W* where the value in *w*_*ij*_ denotes the weight of the directed connection from unit *i* to unit *j*.

### Scores

To evaluate the biological realism of constructed networks, we computed several measures of network activity for both excitatory and inhibitory subpopulations. Networks were evaluated on rate, defined as average spike frequency over the course of each trial. Networks were also evaluated on branching parameter as a measure of network criticality [33]. A branching value of 1 indicates that for every ‘ancestor’ unit that is active, there is an equal number of ‘descendant’ units active at the next time step. On average, the number of units active over the course of a trial in a critical network stays constant. Networks were further evaluated on their level of synchrony, since biological networks display asynchronous activity. In order to evaluate synchrony rapidly enough to make grid search feasible, a heuristic for synchrony was computed as the variance of mean signal normalized by the mean variance of each neuron. A threshold of 0.5 was considered appropriate for network asynchrony. The threshold was evaluated empirically by examining rasters for Poisson spiking neurons with variable coupling, where coupling was change to rate parameter by connected neurons spiking. Branching, was mathematically defined as:

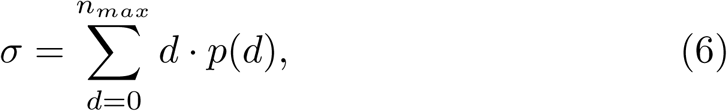

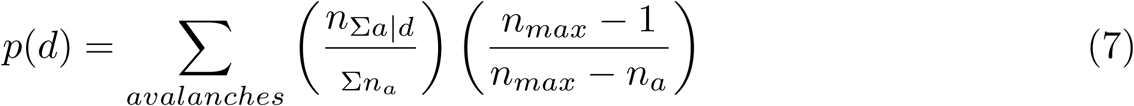

where *σ* is the branching parameter, *d* is the number of descendants, *n*_*max*_ is the maximum number of active neurons, *n*_*a*_ is the number of ancestor neurons, *n*_*d*_ is the number of descendant neurons, *n*_Σ*a*|*d*_ is the number of ancestor neurons in all avalanche events that involved d descendants, and *n*_Σ*a*_ is the total number of neurons involved in avalanches. The branching parameter describes the network as a whole; it cannot be calculated for isolated units. For a given simulation, we calculated network branching at discrete, sequential time steps throughout. We used the same temporal resolution (5 ms) as used for determining the functional graph; all spikes at time *t* are ancestors, and all spikes from *t* + 5 to *t* + 20 ms are descendants. We then averaged the network branching parameter across all time steps to get the overall branching score for that simulation.

For the sake of computational efficiency during grid search, synchrony was defined as the variance of the mean voltage divided by the mean of voltage variances. To evaluate the accuracy of this rapid measure, we compared its results with pairwise Van Rossum spike distance [34, 35] (S1 Fig). We used Van Rossum spike distance as our measure for each simulation’s synchrony score outside of the initial grid search. In this way we are able to generate a multitude of network topologies that produce naturalistic spiking activity.

### Triplet Motifs

The clustering coefficients for the four triplet motifs are calculated in the following manner [43]:

Let *t*_*i*_ denote the actual number of triplets of a motif type in the neighborhood of unit *i*, and *T*_*i*_ denote the maximum number of such triplets that unit i could form. We will build intuition by beginning with the case of a binary directed graph, or an unweighted connectivity matrix. Let *A* represent this graph, with *a*_*ij*_ = 1 indicating the presence of a directed connection from node *i* to node *j*. Raising the matrix *A* to the nth power yields the number of paths of length *n* between nodes *i* and *j*.

Let us first consider the cycle motif; in order for unit *i* to participate in a cycle, it must have an edge directed to a second unit, that second unit must have an edge directed to a third unit, and that third unit must have an edge pointing back to unit *i*. The path length is 3, and it both begins and ends at unit *i*. Thus we calculate *A*^3^ and extract the values along the diagonal, or 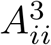. This gives the number of actual cycle motifs unit *i* forms.

Counts of the three isomorphic motifs are calculated in a similar way, but they require the additional involvement of *A*^*T*^. Taking the transpose of graph *A* reverses the directionality, so that connections from *i* to *j* are now those from *j* to *i*. We would like to trace a path of length 3 from i back to i to form an isomorphic triangle, but exactly one of the steps must be against the true direction of that edge (Fig 3). Beginning with a middleman reference node, the first step is ‘with the flow’, the second step is invariably ‘against the flow’, and the final step back to i is again ‘with the flow’. Therefore *AA*^*T*^*A*_*ii*_ gives the number of actual middleman motifs unit i forms. Since fan-in and fan-out motifs are isomorphic to middleman by rotation, we simply rotate which step is ‘against the flow’ to yield the count of fan-in and fan-out motifs. The number of actual fan-in motifs unit i forms is 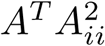, and the number of fan-out motifs is 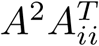.

Now that we can calculate the actual counts, the possible counts of each motif *T*_*i*_ are easily intuited as a combinatorics problem. Let us begin again with the cycle motif. To form a cycle, node i requires one edge directed towards it and one edge directed away from it. The number of possible pairs of in and out edges from node *i* is calculated by multiplying the out-degree of node *i* with the in-degree of node *i*. In-degree and out-degree refer simply to the number of edges that are directed in or out of a given node. Some edges may be bidirectional - these cannot be part of a true cycle motif. The number of bidirectional edges is subtracted from the product of in- and out-degrees. The final *T*_*i*_ for the cycle motif is

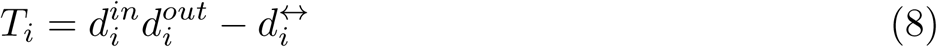

The Ti for middleman is in fact equal to that for cycle, since forming a middleman has the same requirements - one edge directed inward paired with one edge directed outward.

A fan-in motif requires two edges directed in towards the reference node. There are 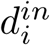 number of choices for the first inward edge. Once that choice has been made, there are 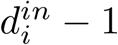 choices remaining for the second inward edge. Thus we multiply the two to yield *T*_*i*_ for the fan-in motif.

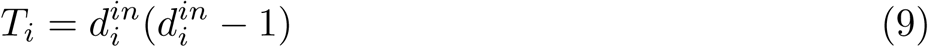

Fan-out is similar - we simply substitute in-degrees with out-degrees since a fan-out motif requires two edges directed out from the reference node. Ti for the fan-out motif is thus

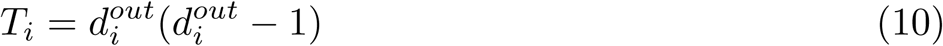

Now that we have both the actual and possible counts for each motif type, the triplet clustering coefficients of node i are simply their ratios. That is,

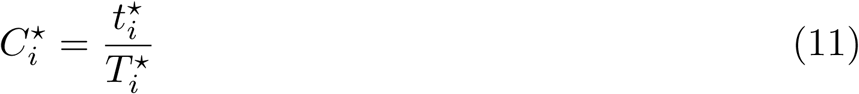

If we were interested in binary graphs, we would end here. However, our graphs of interest have weights associated with each directed edge. There are multiple ways to account for edge weights when calculating clustering coefficients. One way is to consider only the weights of the two edges that are incident to reference node *i*. Alternatively, the weights of all three edges in a triplet can be taken into consideration. The latter is the chosen method, since we desire a measure of central tendency. The total contribution of a triplet to the clustering coefficient is thus the geometric mean of its weights.

Let *W* denote our weighted directed graph. For a triplet in this graph with edge weights *w*_*ij*_, *w*_*ih*_, and *w*_*jh*_, the geometric mean is 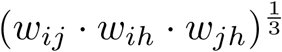. We can extend this to the entire graph by, Instead of using a binary graph as matrix *A* in the calculation of *t*_*i*_, using 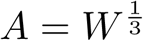, which is the matrix that results from taking the cubic root of every entry in *W*. We also note that this formulation is invariant to the choice of reference node in a triplet. Incorporating weights only modifies the value of *t*_*i*_. It remains a measure of the actual triplets present - instead of counts it is now a weighted measure. The denominator *T*_*i*_ still refers to maximum possible counts. It follows that the clustering coefficient for node *i* can only be 1 (maximum) when its neighborhood truly contains all triplets that could possibly be formed and every edge in each triplet is at unit (maximum) weight. The complete clustering coefficient formulas of weighted directed graphs are given in 3.

### Active Subgraphs

For any small span of time in a trial, only a subset of all units in the graph will be active. The subset of units which spike in some defined time window form the active subgraph for that time window. We binned spikes into a temporal resolution of 10 ms, so that each complete 1-second simulation resulted in 99 time bins. For each time bin t we defined an active subgraph. If a unit spiked within time bin t, that unit will be part of the active subgraph for time bin t. All units which did not spike within that particular time bin are not included in that particular active subgraph. Since there are 99 time bins for a complete 1-second simulation, there are also 99 active subgraphs in sequence. Edge weights between units in an active subgraph are equal to those from the corresponding edges (between active units) in the ground truth synaptic graph.

### Functional Subgraphs

We calculated motifs in the underlying synaptic graphs and found that all four clustering coefficients were equivalent when averaged across each graph, as expected.

To apply motif analysis to activity, we needed to infer functional graphs from spiking activity to summarize network dynamics. Directed edge weights in a functional graph represent the likelihood of a functional relationship in the activity between every pair of units.

We used mutual information (MI) to infer functional graphs from all spikes across the course of a trial, regardless of the trial’s duration (complete or truncated). This results in a single functional graph for each trial. We chose to perform functional inference using the full spike set because this yields functional graphs with higher fidelity and greater sparsity.

The MI method we used is the confluent mutual information between spikes. At a conceptual level, an edge inferred from unit i to unit j using confluent MI means that unit j tends to spike either in the same time bin or one time bin after unit i spikes. Since spikes are binned at 10ms resolution, this method encompasses a delay of 0 to 20 ms. This delay is appropriate because we found that presynaptic spikes yielded a maximal response from all postsynaptic neurons at a delay of 5 to 20 ms.

Mathematically, we defined an indicator function on the spike train of neuron *j, s*(*j*) evaluating to 1 in the case where there is a spike at time *t* or *t* + 1, an indicator function on the spike train of neuron *i, t*(*j*), evaluating to 1 in the case where there is a spike at time *t*, and considered the mutual information between them. The resulting networks were further processed by removing weights corresponding to neurons with negative pairwise correlations. Networks were then re-expressed to minimize skewness, and background signals were removed by accounting for background signal and considering weights as the residual resulting from linear regression on background strength. Finally we considered the z-normalized residual graph to account for heteroskedasticity [56]. This yields weighted values, for which we establish 0 as a threshold. All positive normed residual MI values are included in the full functional graph.

### Recruitment Graphs

The recruitment graph represents both the activity and the underlying connections of a network. A recruitment graph is defined separately for each 10 ms time bin of a given trial, thus yielding a temporal sequence of graphs. Each graph is calculated as the intersection of the functional graph, which is unique to every trial, and the active subgraph, which is unique to every 10 ms time bin. All edges in the recruitment graph come from underlying synaptic wiring, contained in the active subgraph, while edge weight values come from the inferred functional graph. In other words, for all edges *i* → *j* where *w*_*functional,ij*_ *>* 0 in the confluent MI functional graph and *w*_*synaptic,ij*_ *>* 0 in the active subgraph of time bin *t*, the edge in the recruitment graph for time bin t takes on the value of *w*_*functional,ij*_. All other recruitment graph edges have value 0.

Just like the sequence of active subgraphs, there are 99 sequential recruitment graphs at 10 ms temporal resolution for every complete 1-second simulation trial. Triplet clustering coefficients were calculated for every unit on each 10 ms recruitment graph, then averaged across the population to yield the whole-network clustering coefficients for that 10 ms time window. These methods allow us to observe how motif clustering changes in the recruitment graphs across time.

### Clustering Propensity

Networks may have very different connection densities, which would impact motif clustering coefficients. This is especially true as the active subnetwork changes in time, and for ER networks in comparison to clustered model graphs. We therefore used weighted and unweighted clustering propensity which enables meaningful comparisons between networks with different connection densities.

Our measure of weighted propensity begins with calculating the triplet motif clustering coefficients for each unit in the recruitment graph of every time bin. Then, for each time bin t we generate ten simulated graphs. These graphs have the same edges as the original recruitment graph at time *t*, with edge weights randomly sampled from the underlying distribution of functional edge weights. Motif clustering coefficients are calculated for units in each of the simulated graphs, then averaged for each unit and each motif type. The clustering coefficients of the units in the original graph at time *t* are normalized by the average of the ten simulated graphs’ clustering coefficients, yielding the unit-wise clustering propensity at time *t* for each triplet motif. We used these values to perform all unit-wise motif analysis. In order to examine motifs at a whole-network level, for each motif type at time *t* we average across all units with nonzero clustering propensity values for that motif type.

Unweighted propensity is calculated similarly, considering the functional networks’ unweighted directed clustering to that expected in both an ER graph as well as a small world graph. Thus, in addition to controlling for density, weighted propensity also measures the extent to which the specific edge weights in the recruitment graph impact triplet motif clustering, while unweighted propensity measures the same for specific structure of the recruitment graph. A weighted propensity of 1 indicates that specific edge weights play a negligible role in clustering, since random edge weights would still yield the same clustering coefficients, while an unweighted propensity of 1 indicates that the specific structure of the network is not important for clustering.

### Erdős-Renyi Graph Simulations

Unclustered ER Graph simulations were done from population consisting of 1000 excitatory neurons, 200 inhibitory neurons and 50 poisson input neurons. These populations were connected with *p*_*ee*_ = .2, *p*_*ii*_ = .3, *p*_*ie*_ = .25 and *p*_*ei*_ = .35. Synaptic weights relative to leak conductance were drawn from a log normal distribution (mean=.60, variance = .11), with i to e connections scaled up 50% [19].

### Overlap Index

Overlap Index was used to measure the degree of overlap between two probability distributions. It is defined as 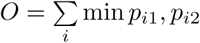, where *i* is histogram bin index. If two distributions do not overlap at all they will have an overlap index of 0, if they are identical they will have an overlap index of 1.

### Probability Vectors

To quantify the cyclic transitions between relative prominence of motifs over time, we examined the dominant motif of the network for a given recruitment graph. We define the dominant motif of a graph as the maximum of the demeaned propensities for middleman, fan-in, and fan-out. We consider the demeaned values of each motif in order to account for the different relative magnitudes of motifs without affecting scaling in the cycle structure. Examining first order probabilities, which is the probability of a motif dominating a recruitment graph in a given run, on the time series of recruitment graphs from completed and truncated runs shows that there is a significant difference between the distributions defining these values across each type of run.

To further characterize the transitions between different dominant motifs we fit a Markov model to the series of dominant motifs across recruitment graphs in both truncated and completed networks. We again find a significant different (*p <* .001 for all markov parameters). This all suggests that the failure to propagate found in some networks is tied to the inability to recruit the cyclic structure that we find to be a hallmark of successful propagation.

### Statistical Testing: Two-Sample Chi Squared Test

P values for comparisons between distributions of different types of network activity were done by two sample chi square test.

## Supporting information

**S1 Fig.**
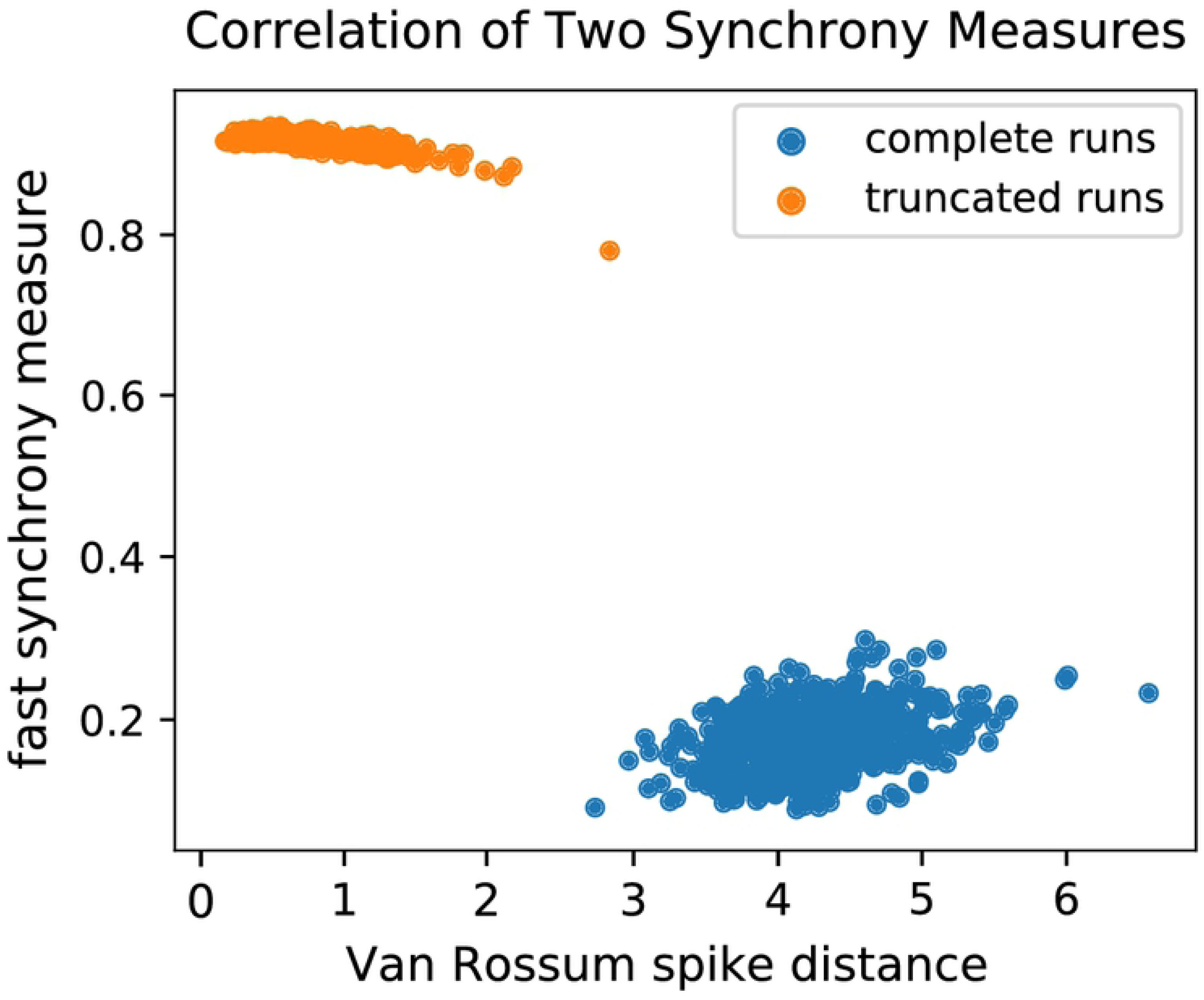
Synchrony Measure Comparison. A common measurement for synchrony is the Van Rossum spike distance [34, 35]. Greater distances between spike pairs indicate more asynchronous dynamics. However, for the sake of speed during grid search, we used a rapid measure of synchrony (see Methods). We observe a strong correlation between the two measures when we used both to examine the final set of runs that we used for analysis..

**S1 Table.** Neuron Parameters. Parameters used for simulation of adaptive exponential integrate and fire neurons.

| Parameter | Name | Value |
| --- | --- | --- |
| $C$ | membrane capacitance | 281 pF |
| $g_L$ | leak conductance | 30 nS |
| $E_L$ | leak reversal potential | -70.6 mV |
| $E_E$ | excitatory reversal potential | 0 mV |
| $E_I$ | inhibitory reversal potential | -75 mV |
| $V_T$ | spike threshold | -50.4 mV |
| $\Delta_T$ | slope factor | 2 mV |
| $\tau_w$ | adaptation time constant | 144 ms |
| $\tau_e$ | excitatory time constant | 10 ms |
| $\tau_i$ | inhibitory time constant | 3 ms |
| $\tau_p$ | poisson time constant | 3ms |
| $a$ | subthreshold adaptation | 4 nS |
| $b$ | spike triggered adaptation | .0805 nA |

## Acknowledgments

We thank Elizabeth de Laittre, Maayan Levy, Graham Smith, and Barbara Peysakhovich for helpful comments on our manuscript.

